# Resistance Rate Distribution of MDR-TB Among Pulmonary Tuberculosis Patients Attending Nnamdi Azikiwe University Teaching Hospital Nnewi and ST Patrick’S Hospital Mile 4 Abakiliki in Southeast Nigeria

**DOI:** 10.1101/570416

**Authors:** Chinenye Esther Okoro, Confort Nnenna Akujobi, Iniekong Philip Udoh, Stellamaris Ojiuzor Ibhawaegbele, Charles Ikechukwu Ezema, Uchechukwu Anthonia Ezeugwu, Ogechukwu Calista Dozie-Nwakile, Aaron Chukwuemeka Okpe

## Abstract

Tuberculosis, one of the oldest recorded human afflictions, is still one of the biggest killers among the infectious diseases, despite the worldwide use of a live attenuated vaccine and several antibiotics. This study was designed to assess the resistance rate distribution of MDR-TB among pulmonary tuberculosis patients attending Nnamdi Azikiewe University Teaching Hospital (NAUTH) Nnewi and St Patrick’s Hospital Mile 4 Abakaliki in the Southeast Nigeria. Patients with persistent cough for over two weeks were screened by Ziehl-Neelsen (ZN) technique for the presence of acid fast bacilli (AFB) in their sputum and a total of 103 patients with AFB positive sputum samples were recruited. The positive sputum samples were subjected to Xpert MTB/RIF assay (GeneXpert®, Cepheid USA) and culture on Lowestein Jensen medium for 42days at 37°C. Drug susceptibility testing was done on the isolates using the nitrate reduction assay (NRA). Xpert MTB/RIF assay detected MTB in 83(80.6%) samples out of which 45(67.2%) were rifampicin resistant. Sixty-seven (80.7%) of the isolates were resistant to at least one of the first-line drugs. Primary resistance was 91% while 19.4%, 35.8%, 22.4% and 22.4% of the isolates were resistant to one, two, three and four drugs respectively. Isoniazid had the highest rate of resistance (57.8%) while Ethambutol had the least (34.9%) and 30(44.8%) of the resistant isolates were MDR. Smoking (P=.002), gender (P=.002) and history of TB treatment (P=.012) were significantly associated with drug resistance. Educational status was significantly associated with MDR-TB (P=.020). NAUTH and St Patrick’s hospital had MDR-TB rates of 38.9% and 46.9% respectively. The findings of this study indicate high prevalence of MDR-TB among patients with pulmonary TB in the study sites and this portrays a menace to adequate TB control. Prompt diagnosis of TB, adequate patient compliance to therapy and increased awareness and mass education is recommended

## INTRODUCTION

The discovery of anti-tuberculosis drugs in the 1940s followed by combination chemotherapy made tuberculosis a curable disease. In the developed countries, effective treatment and surveillance reduced tuberculosis dramatically with high hopes of total eradication [1, 2]. However, in the 1980s, it was realized that tuberculosis had not only ceased to decline in the developed countries, notably the USA, but was actually increasing, particularly in major cities [2]. It was also soon realized that the disease was out of control and increasing at an alarming rate across most of the poorest regions of the world especially Africa due to HIV/AIDS [1, 3]. Despite aggressive international efforts, tuberculosis remains a leading infectious cause of death, with an estimated 8.6 million incident cases per year. In 2012, an estimated 1.3 million people died from the disease. These death rates, however, only partially depict the global TB threat; more than 80% of TB patients are in the economically productive age of 15 to 49 years [4].

Interestingly, global tuberculosis control efforts have been threatened by the emergence of multidrug resistant tuberculosis (MDR-TB). It is the strains of *Mycobacterium tuberculosis* which show high level resistance to both isoniazid and rifampicin, with or without resistance to other anti TB drugs [4]. Alarmingly, MDR-TB is estimated to cause 4% of new tuberculosis cases in the developing world. Patients infected with MDR strains are not only difficult to cure but also more likely to remain sources of infection for a longer period of time than those with drug susceptible organisms. MDR-TB requires longer duration of treatment (up to 2 years) to achieve cure, in comparison with 6 month treatment for drug susceptible TB, lower cure rates and even higher default rates, not minding the expensive cost of treatment [5].

Remarkably, due to the increasing prevalence, MDR-TB is now subdivided into basic MDR-TB, with resistance only to rifampicin and isoniazid, and extensive drug resistant TB (XDR-TB), with a similar resistance pattern but with resistance to one or more additional first and/or second line drugs. Various perturbations in the individual drug target genes are responsible for the genesis of anti-TB drugs resistance. Rifampicin resistance has been shown to be caused by a change in the β-subunit of DNA dependent RNA polymerase, which is encoded by the *rpoβ* gene and more than 95% of rifampicin resistant strains are associated with mutations within an 81-base pair region of the *rpoβ* gene, which is termed rifampicin resistance determinant region [6, 7, 8]. On the contrary, resistance to isoniazid is due to mutations at one of two main sites, in either the *katG* or *inhA* genes [9, 10]. It is also noted that these mutations are not directly connected, and so separate mutations are required for organisms to change from a drug susceptible isolate to MDR-TB. Furthermore, rifampicin resistance has been considered to be a surrogate marker for checking multidrug resistance in clinical isolates of *M*. *tuberculosis* since rifampicin resistance is often accompanied by resistance to isoniazid [7, 8]. Drug resistance in *M. tuberculosis* occurs by random, single step, spontaneous mutation at a low but predictable frequency, in large bacterial populations. The accurate diagnosis of MDR-TB requires a positive culture of *M. tuberculosis* and drug susceptibility testing. The use of genotypic analysis of rpoβ for Rif resistance in evaluating the public health threat of *Mycobacterium tuberculosis* is controversial due to the fact that misdiagnosing of patients as MDR-TB when they are only Rif mono-resistant would lead to inappropriate second line treatment in a world of limited second line armamentarium [11].

Due to the burden in diagnosing MDR-TB in pulmonary tuberculosis patients resulting from poor facilities in Nigeria, this study therefore assessed the resistance rate distribution of MDR-TB among pulmonary tuberculosis patients attending Nnamdi Azikiewe University Teaching Hospital Nnewi and St Patrick’s Hospital Mile 4 Abakaliki in the Southeast Nigeria.

## MATERIALS AND METHODS

### Study Area

This study was conducted at Nnamdi Azikiwe University Teaching Hospital (NAUTH), Nnewi and St Patrick’s Hospital, Mile 4 Abakaliki. Nnewi is the second largest city in Anambra State and is home to nearly 388,805 residents. Abakaliki is the capital of Ebonyi state and has a population of 149,683 persons [12]. NAUTH is a tertiary health institution and serves as a site for treatment and management of both TB and HIV patients. It is also a referral centre for both cases. St Patrick’s Hospital, Mile 4 Abakaliki is a faith-based health facility and offers both antiretroviral therapy and TB care to patients.

### Sample Size

Minimum sample size was calculated using the formula stated by [13] and a total of 103 sputum smear positive AFB samples were collected for the study.

### Ethical Approval

Ethical approval for this study was obtained from NAUTH research and ethics committee. Consent was obtained from each participant and participants’ confidentiality was maintained throughout the study. Participants received no financial motivation for their involvement in the study. Participants were free to withdraw from the study at any point and their withdrawal would not affect their treatment. This study was conducted between January 2015 and September 2016

### Sample collection and analysis

About 2mls of venous blood sample from smear positive AFB participants’ was collected in plain tube, allowed to clot and the serum separated and screened for presence of HIV-1/2 antibodies using serial algorithm method. Determine®, Unigold® and Stat-pak® HIV test kits were used according to manufacturer’s instruction (Determine^®^ is manufactured by Alere Medical Co., Ltd Japan while Unigold^®^ and StatPak^®^ are manufactured by Trinity Biotech PLC, Ireland and CHEMBIO Diagnostic Systems Inc New York, USA respectively) [14].

Consenting, eligible participants were screened for presence of AFB in their sputum. Two sputum samples (spot and early morning) were collected in sterile screw-cap universal containers from each participant on 2 consecutive days and stained by Ziehl-Neelsen’s method.

Progressively, early morning mucoid or mucopurulent sputum specimen was collected from each participant with smear positive AFB test result into a sterile screw-cap universal bottle. The specimen was then stored in the refrigerator until transported to the TB reference laboratory of Dr Lawrence Henshaw Memorial Hospital (DLHMH) in Calabar, Cross River State. Transport was done within 72hrs of collection.

After appropriate sample preparation, two Lowestein Jensen (LJ) medium slants were cultured for each sample. Tubes were loosely capped and incubated as such at 37°C for one week in a slanted position to ensure even distribution and absorption of inoculum. After 1 week, tubes were incubated upright for up to 6 weeks and the caps tightened. An in-house strain H37RV and an uninoculated tube were used as positive and negative control respectively as previously reported by [15].

After Colonies was confirmed by Ziehl-Neelsen (ZN) staining for acid-fastness, niacin test was carried out on each inoculated and control tubes. The formation of a yellow colour was interpreted as positive reaction; absence of colour was regarded as negative reaction for production of Niacin. Catalase test, p-Nitrobenzoic Acid (PNB) and TB Ag MPT64 Rapid Test was carried out in this study and *M. tuberculosis* identification was based on its slow growth rate, no pigmentation, no growth on Lowestein Jensen (LJ) medium containing p-nitrobenzoic acid, niacin production, catalase negative at 68°C and positive Ag MPT 64 test.

Drug susceptibility testing (DST) was carried out on all confirmed *M. tuberculosis* colonies and nitrate reduction assay (NRA) method was used [16].

GeneXpert MTB/RIF assay for detection of Rifampicin Resistance was carried out on the sputum samples of the participants. Sputum sediments were mixed with sample buffer in a ratio of 1:3 in a screw cap tube and screwed tightly. The tube was vortexed for 20 seconds. Sample was incubated at room temperature for 10mins. After 10mins the sample was vortexed again for 20 seconds and incubated at room temperature for 5mins. After incubation, 2ml of sample was inoculated into the genexpert cartridge. Cartridge was scanned into the GeneXpert machine (Cepheid USA) and allowed to run for 2hrs. After 2hrs the test result was read off the screen of the GeneXpert machine monitor.

### Data Analysis

Data was statistically analyzed using statistical package for social sciences SSPS for windows version 20.0 software. A standard questionnaire was completed for each recruited patient to collect demographic parameters. Frequencies were calculated as percentages. Comparison of categorical variables and significance testing was done with χ^2^ test. P-value of less than 0.05(P<0.05) was considered statistical significant.

## RESULTS

Out of the 103 AFB positive sputum samples collected 83(80.6%) showed culture positive isolates. Sixty-one (61) of the isolates were from St Patrick’s Hospital, Mile 4 Abakaliki while 30 were from NAUTH giving a TB prevalence rate of 73.50% and 26.50% respectively. Fig 2 showed 80.70% resistance rate of the isolates. NAUTH had resistance rate of 81.80% compared with 80.70% from Mile 4 Abakaliki.

**Fig 1:**
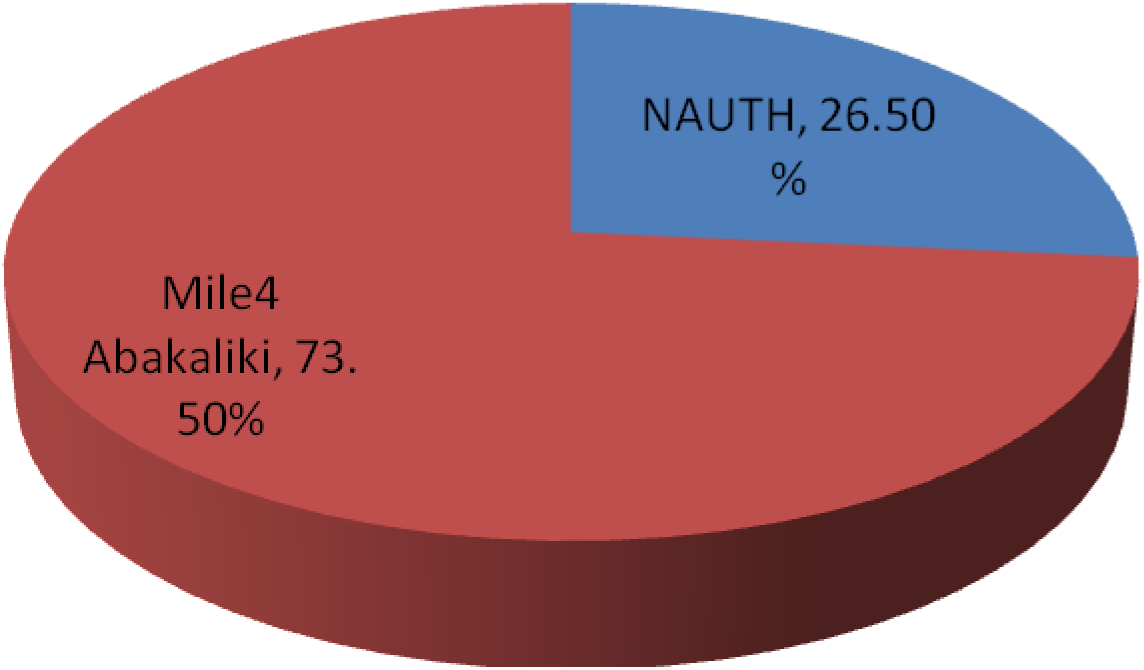
Culture Positivity based on Site.

**Fig 2:**
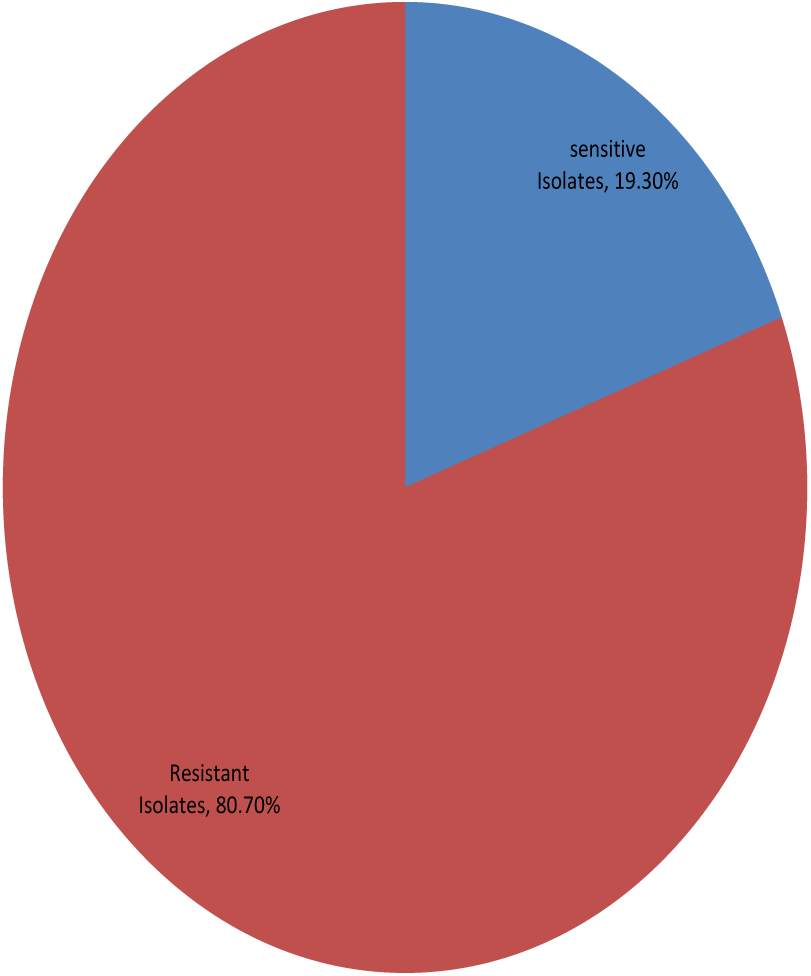
Resistance Rate of Isolates in the Study.

Table 1 showed no statistically significant difference among the age groups, though age group 18-25years and 26-35years showed high resistance rate of (91.3%) and (77.8%) respectively. Gender showed statistically significant association with resistance to first-line anti-TB drugs. Employment status, educational status, residence and marital status showed no significant difference. In table 2, history of smoking and previous TB treatment was found to be statistically significant for drug resistance.

**Table 1:**
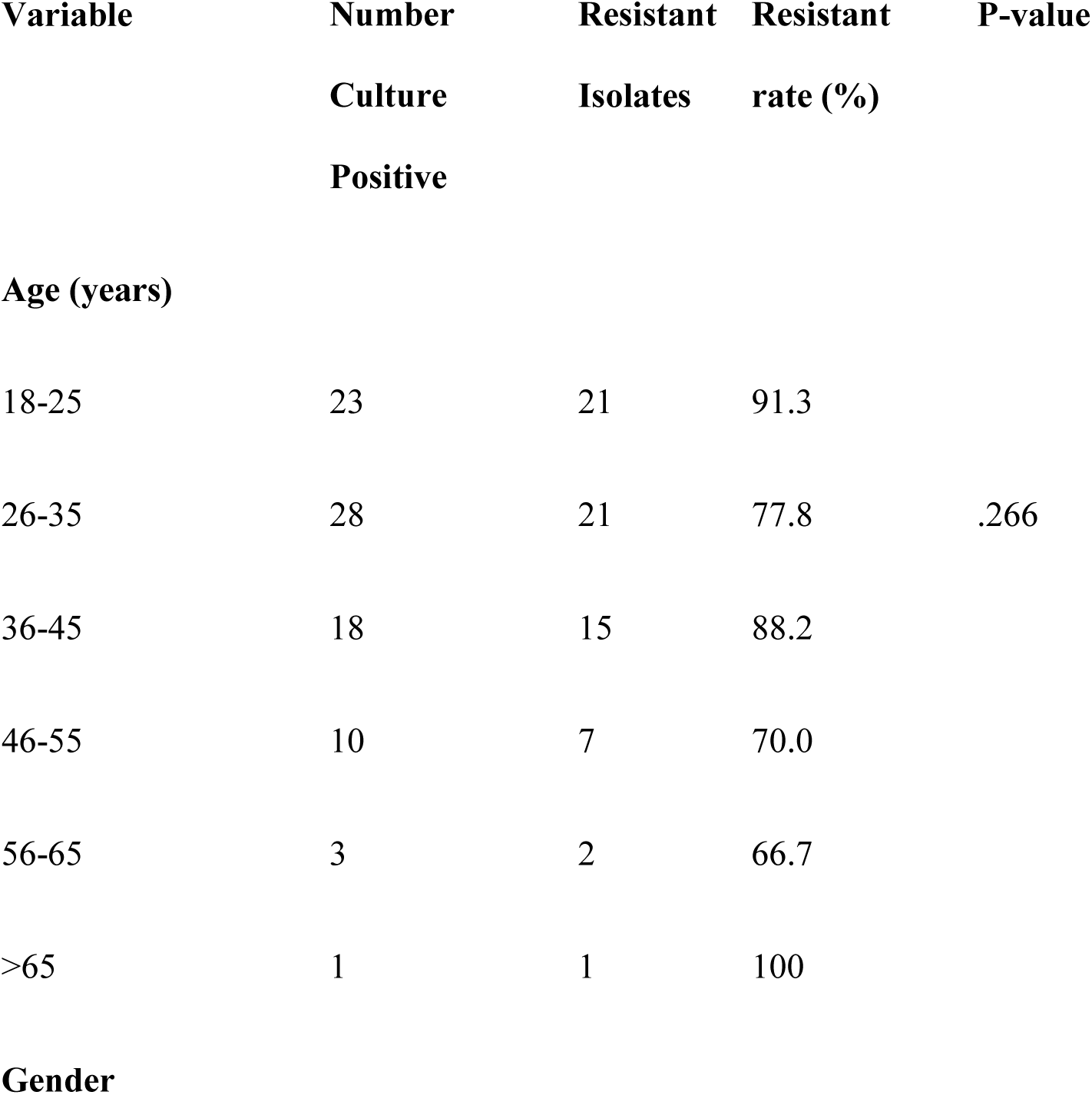
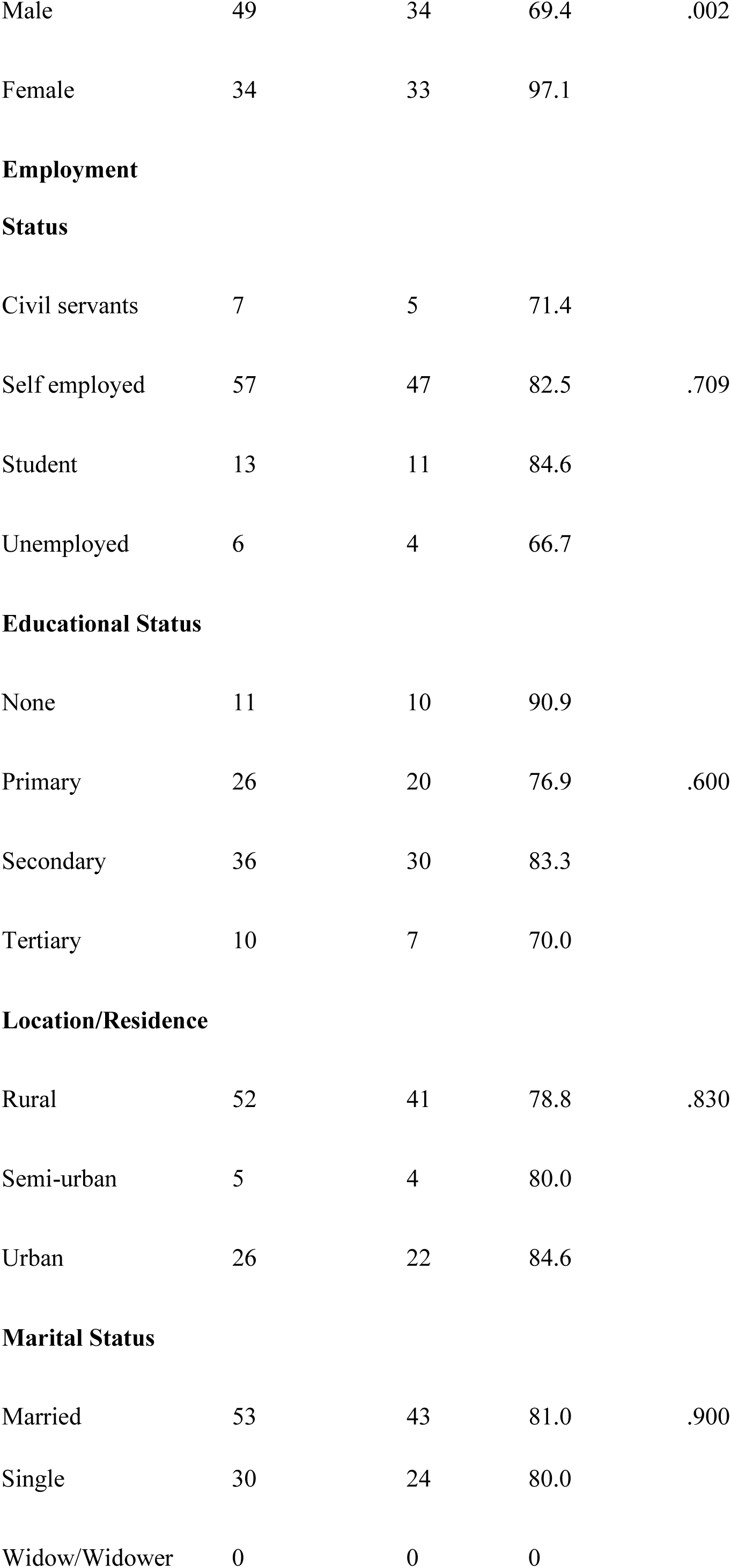
Resistance Rates of Isolates with respect to Demographic Factors.

**Table 2:**
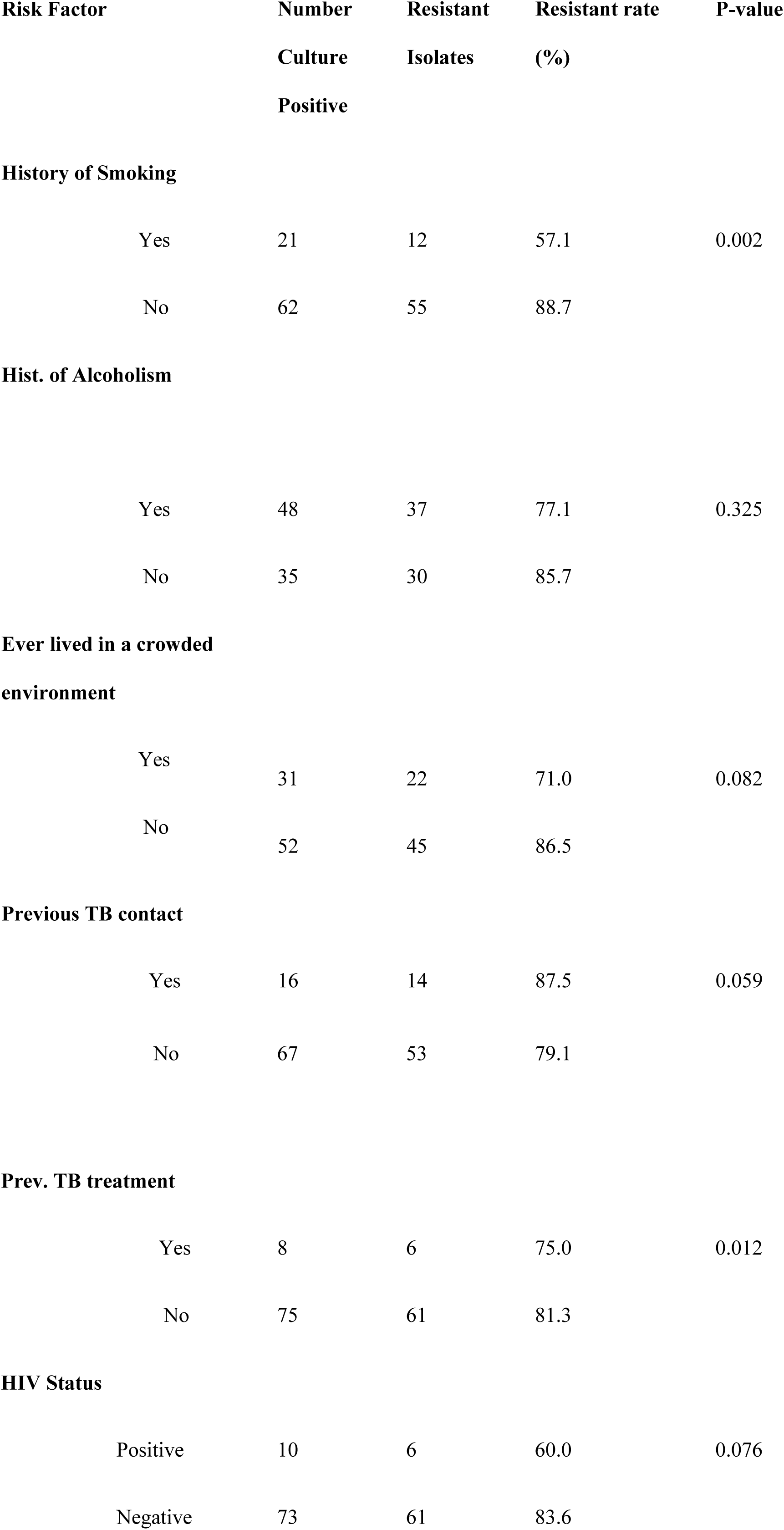
Assessment of Some Risk Factors with respect to Drug Resistance.

| Risk Factor | Number | Resistant | Resistant rate | P-value |
| --- | --- | --- | --- | --- |
|  | Culture | Isolates | (%) |  |
|  | Positive |  |  |  |
| History of Smoking |  |  |  |  |
| Yes | 21 | 12 | 57.1 | 0.002 |
| No | 62 | 55 | 88.7 |  |
| Hist. of Alcoholism |  |  |  |  |
| Yes | 48 | 37 | 77.1 | 0.325 |
| No | 35 | 30 | 85.7 |  |
| Ever lived in a crowded environment |  |  |  |  |
| Yes | 31 | 22 | 71.0 | 0.082 |
| No | 52 | 45 | 86.5 |  |
| Previous TB contact |  |  |  |  |
| Yes | 16 | 14 | 87.5 | 0.059 |

|  |  |  |  |
| --- | --- | --- | --- |
| No | 67 | 53 | 79.1 |
| **Prev. TB treatment** |  |  |  |  |
| Yes | 8 | 6 | 75.0 | 0.012 |

|  |  |  |  |
| --- | --- | --- | --- |
| No | 75 | 61 | 81.3 |
| **HIV Status** |  |  |  |  |
| Positive | 10 | 6 | 60.0 | 0.076 |

|  |  |  |  |
| --- | --- | --- | --- |
| Negative | 73 | 61 | 83.6 |

Fig 4 showed primary and acquired resistance rate prevalence of the study. It shows also that primary resistance (91%) were higher than the acquired resistance (9%). Fig 5 showed the degree of the resistance of the isolate to first line anti-TB drugs 19.4%, 35.8%, 22.4% and 22.4% of the isolates were resistant to one, two, three and four drugs respectively. In fig 6, isoniazid (57.80%) showed the highest resistance rate followed by rifampicin (54.20%), streptomycin (53.00%) and least by ethambutol (34%).

**Fig 3:**
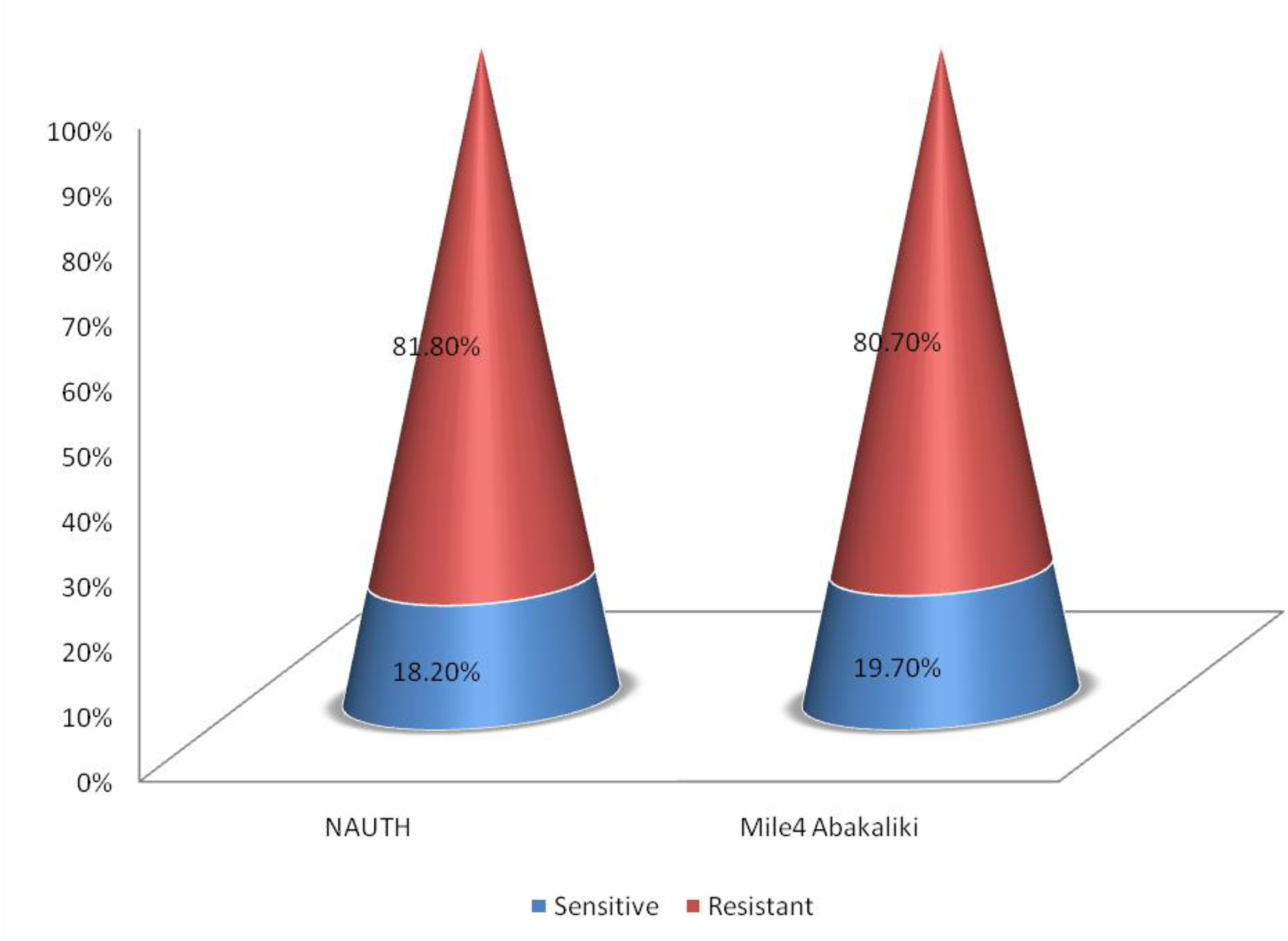
Resistance Rate of Isolates based on Site Abbreviations **NAUTH:** Nnamdi Azikiwe University Teaching Hospital

**Fig 4:**
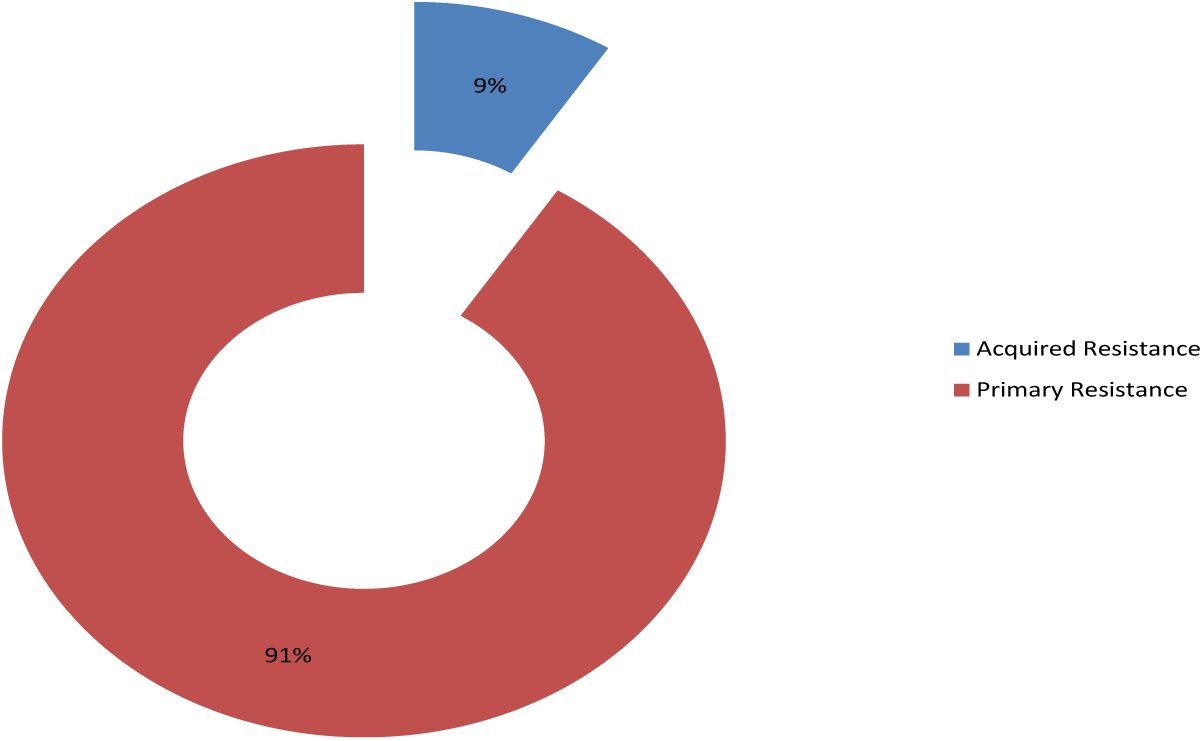
Rate of primary (treatment naïve) resistance in the study.

**Fig 5:**
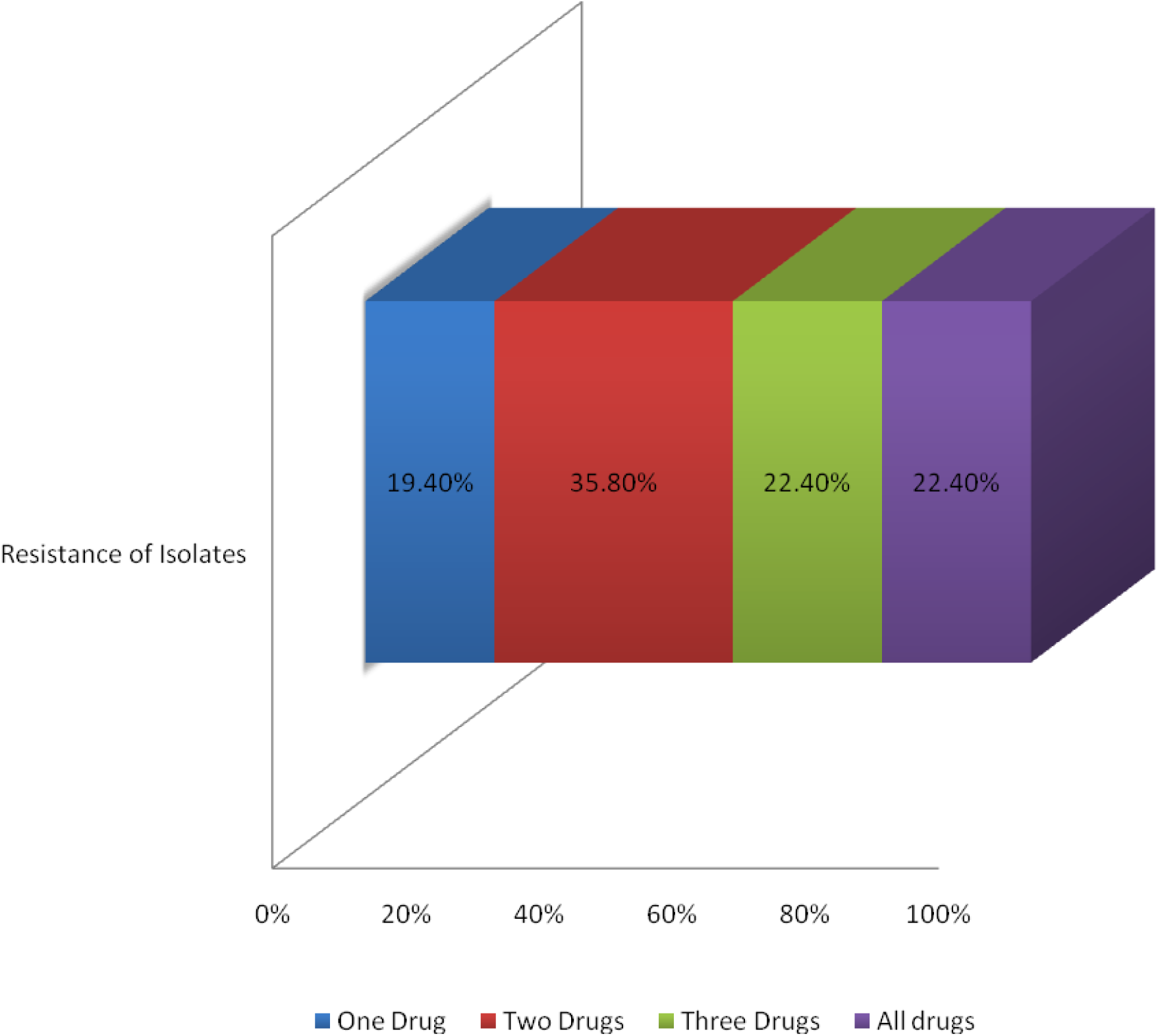
Degree of Resistance of Isolates to First-line Anti-TB Drugs.

**Fig 6:**
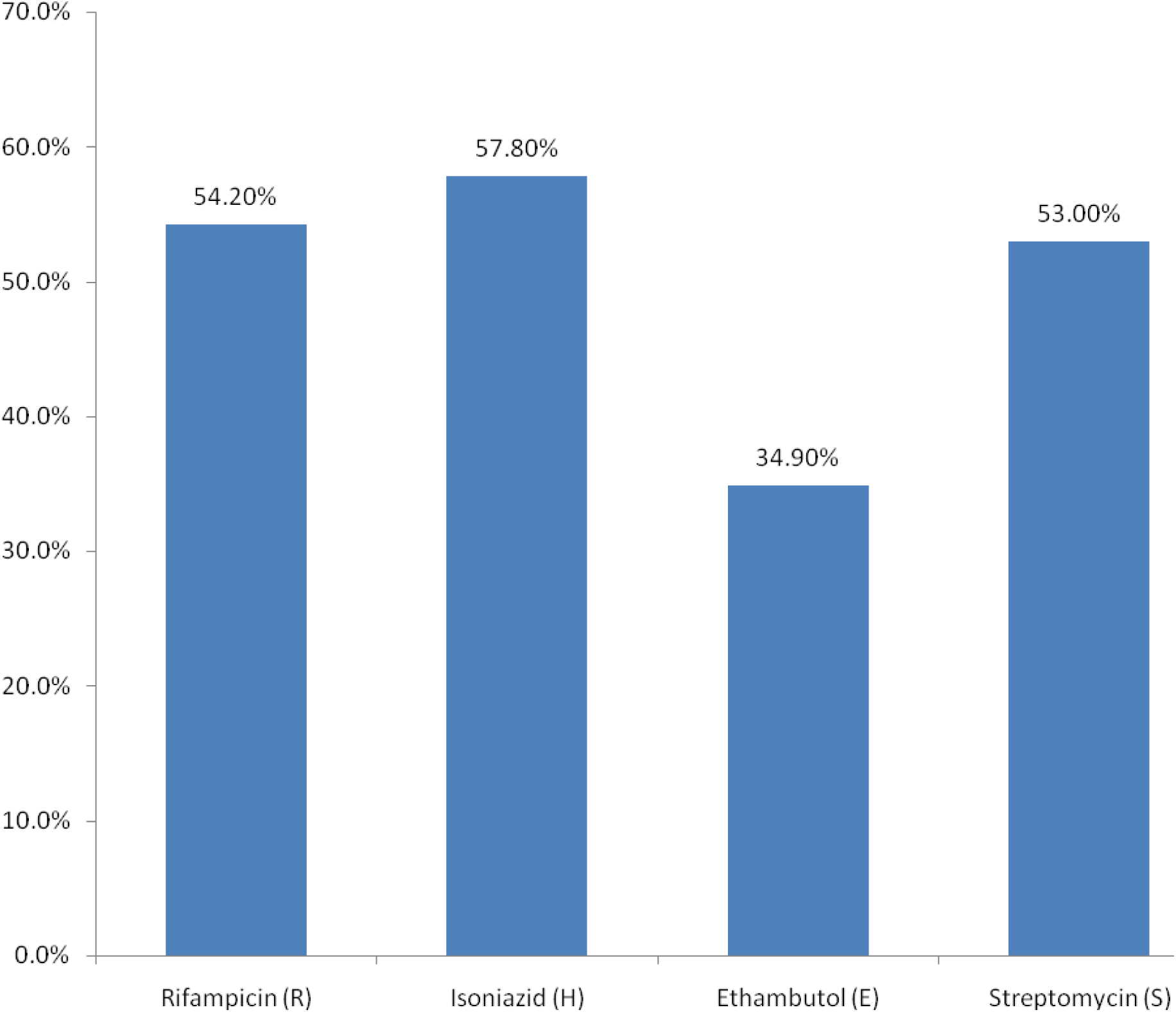
Resistance rate of isolates to individual first-line Anti-TB Drugs.

In table 3, isolates of *M. tuberculosis* exhibited fourteen resistance patterns to the first-line anti-TB drugs with the combination of Streptomycin, Isoniazid, Rifampicin and Ethambutol showing the highest resistance rate (22.4%). However based on site, table 4 showed that isolates from NAUTH exhibited eleven (11) distinct resistance patterns while isolates from Mile 4 Hospital, table 5, exhibited fourteen (14) patterns. Table 6, showed resistance pattern of isolates with respect to treatment history, treatment Naïve (primary treatment) exhibited higher resistance pattern when compared with retreatment. Fig 7 showed high Prevalence of **MDR-TB** (37.50%) with respect to treatment history than retreatment (36.00%).

**Table 3:** Pattern of Resistance of Isolates to First-line Anti-TB Drugs.

| **Drug Pattern** | **No. of Isolates Resistant (%)** |
| --- | --- |
| **Rifampicin only** | 1 (1.5) |
| **Ethambutol only** | 1 (1.5) |
| **Isoniazid only** | 8 (11.9) |
| **Streptomycin only** | 3 (4.5) |
| **Rifampicin and Ethambutol** | 2 (3.0) |
| **Rifampicin and Streptomycin** | 9 (13.4) |
| **Isoniazid and Ethambutol** | 4 (6.0) |
| **Isoniazid and Streptomycin** | 4 (6.0) |
| **Isoniazid and Rifampicin** | 5 (7.5) |
| **Rifampicin, Isoniazid and Ethambutol** | 2 (3.0) |

**Table 4:**
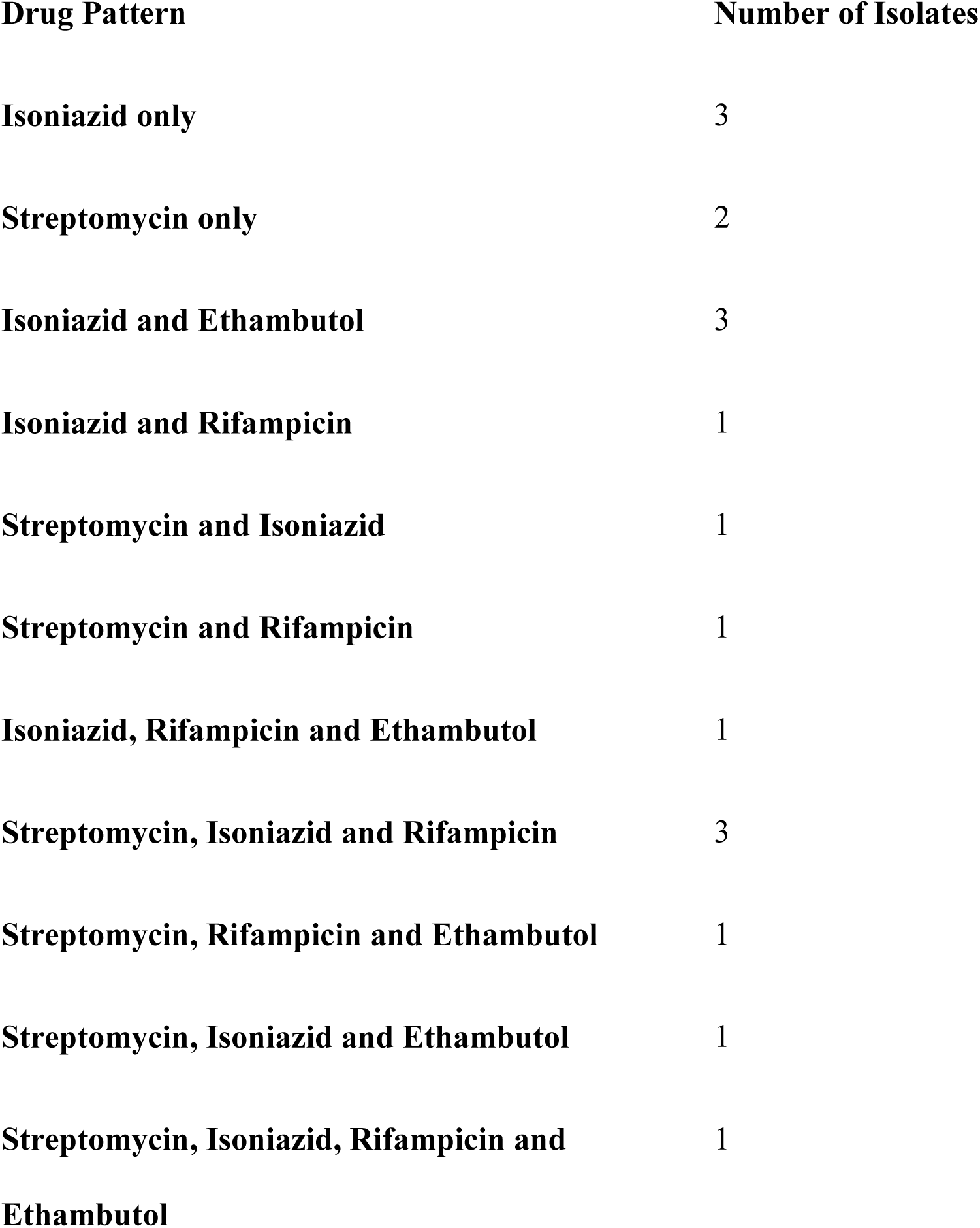
Resistant Pattern of Isolates from NAUTH.

| **Drug Pattern** | **Number of Isolates** |
| --- | --- |
| **Isoniazid only** | 3 |
| **Streptomycin only** | 2 |
| **Isoniazid and Ethambutol** | 3 |
| **Isoniazid and Rifampicin** | 1 |
| **Streptomycin and Isoniazid** | 1 |
| **Streptomycin and Rifampicin** | 1 |
| **Isoniazid, Rifampicin and Ethambutol** | 1 |
| **Streptomycin, Isoniazid and Rifampicin** | 3 |
| **Streptomycin, Rifampicin and Ethambutol** | 1 |
| **Streptomycin, Isoniazid and Ethambutol** | 1 |
| **Streptomycin, Isoniazid, Rifampicin and Ethambutol** | 1 |

**Table 5:** Resistant Pattern of Isolates from Mile 4 Hospital Abakaliki.

| **Drug Pattern** | **No. of Isolates Resistant** |
| --- | --- |
| **Rifampicin only** | 1 |
| **Ethambutol only** | 1 |
| **Isoniazid only** | 5 |
| **Streptomycin only** | 1 |
| **Rifampicin and Ethambutol** | 2 |
| **Rifampicin and Streptomycin** | 8 |
| **Isoniazid and Ethambutol** | 1 |
| **Isoniazid and Streptomycin** | 3 |
| **Isoniazid and Rifampicin** | 4 |
| **Rifampicin, Isoniazid and Ethambutol** | 1 |
| **Streptomycin, Isoniazid and Rifampicin** | 5 |
| **Strptomycin, Rifampicin and Ethambutol** | 2 |
| **Streptomycin, Isoniazid and Ethambutol** | 1 |
| **Streptomycin, Isoniazid, Rifampicin and Ethambutol** | 14 |

**Table 6:** Resistance pattern of Isolates with respect to Treatment History.

| **Drug Pattern** | **No. of Resistant Isolates** |  |
| --- | --- | --- |
|  | **Treatment Naive** | **Retreatment** |
| **Rifampicin only** | 1 | 0 |
| **Ethambutol only** | 1 | 0 |
| **Isoniazid only** | 8 | 0 |
| **Streptomycin only** | 3 | 0 |
| **Rifampicin and Ethambutol** | 1 | 1 |
| **Rifampicin and Streptomycin** | 9 | 0 |
| **Isoniazid and Ethambutol** | 4 | 0 |
| **Isoniazid and Streptomycin** | 2 | 2 |
| **Isoniazid and Rifampicin** | 4 | 1 |
| **Rifampicin, Isoniazid and Ethambutol** | 1 | 1 |
| **Streptomycin, Isoniazid and Rifampicin** | 7 | 1 |
| **Strptomycin, Rifampicin and Ethambutol** | 3 | 0 |
| **Streptomycin, Isoniazid and Ethambutol** | 2 | 0 |
| **Streptomycin, Isoniazid, Rifampicin and Ethambutol** | 15 | 0 |

**Fig 7:**
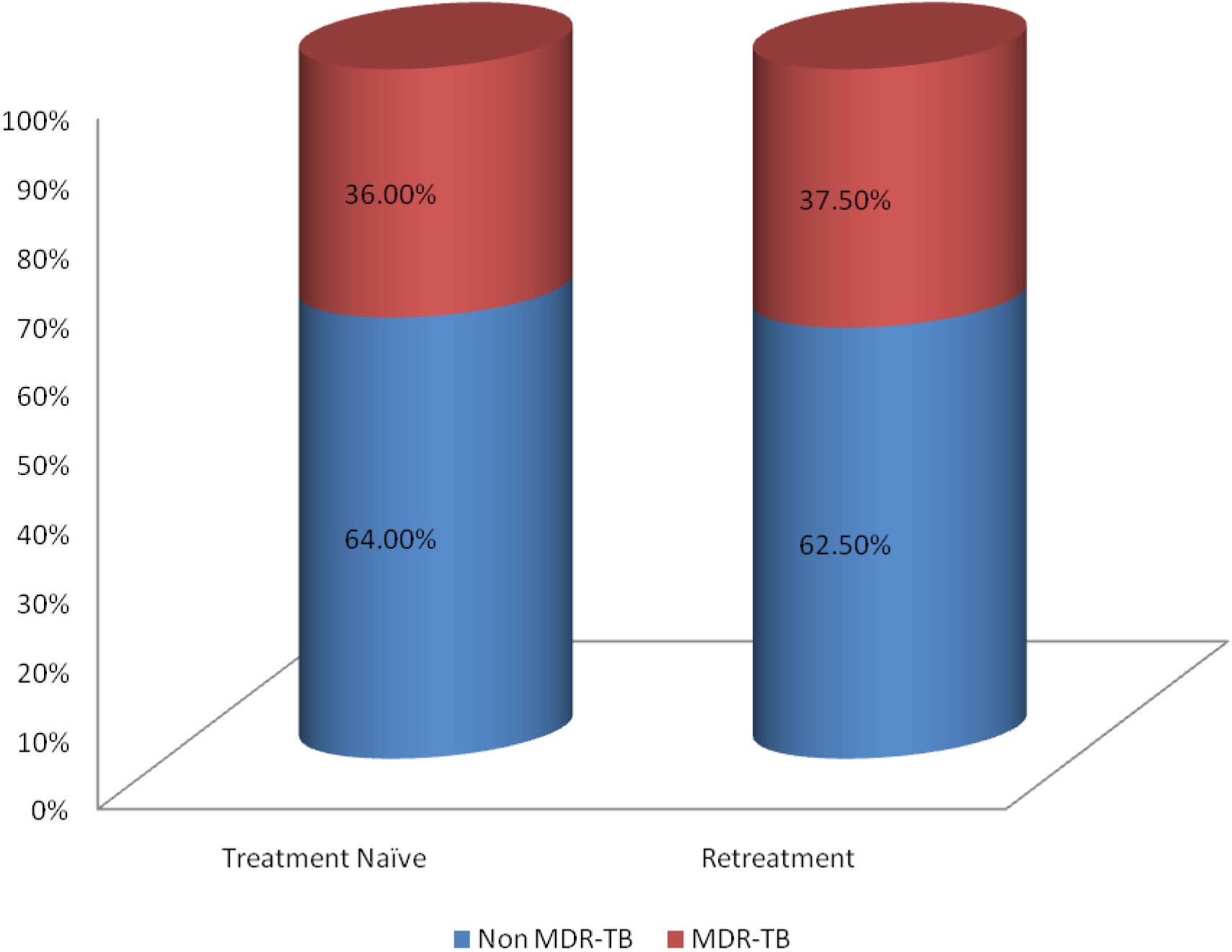
Prevalence of MDR-TB with respect to Treatment History. **Abbreviations**
Non MDR-TB: Non multidrug resistance Tuberculosis
MDR-TB: Multidrug resistance Tuberculosis

## DISCUSSION

Tuberculosis remains the major global health problem which ranked the 9th leading cause of death worldwide and currently, the emergency of MDR-TB is also the main public health problem in both developing and developed countries [17]. Drug resistance in *M. tuberculosis* isolates arises from spontaneous genetic mutations and can be enhanced by poor adherence of patients to anti-TB drugs [18]. Moreover, infection depends both on the bacterial virulence and the inherent microbicidal ability of the alveolar macrophage that ingests it. If the bacillus is able to survive initial defences, it can multiply within the alveolar macrophage [19]. Interestingly, out of a total of 103 acid fast bacilli AFB positive samples in this study 83 (80.6%) yielded *Mycobacterium tuberculosis* isolates on culture, 12(11.7%) were negative for *M. tuberculosis*, while 8(7.8%) were contaminated. Sixty-one (61) of the isolates were from St Patrick’s Hospital, Mile 4 Abakaliki while 30 were from NAUTH giving a TB prevalence rate of 73.50% and 26.50% respectively. The high culture positivity rate in this study agrees with [20] that reported 65.7% culture positivity rate in South-West Nigeria. Also [21] reported that 100 out of 120 sputum samples were positive for *M. tuberculosis* on culture in a study in Calabar. In the contrary, this rate is higher than the 33% culture positivity rate reported by [15] in a study at Nnewi. Also [22] reported a lower rate of 44% in India.

Report has it that drug resistance is mainly due to irregular or improper anti-TB drug use and absence of good, effective national TB control programme which have led to accumulation and multiplication of resistance strains [23, 22]. The high resistance rate of isolates in this study is similarly high in consonance with [24]. They reported that 81.2% of their isolates showed resistance to at least one drug in Georgia. [20] Reported a resistance rate of 62.5% in South-West Nigeria. Also [22] reported 69.7% resistance rate in a study conducted in India. The resistance rate obtained in this study however is higher than the 31% resistance rate reported by [25] in a study at Abuja and this could be due to geographical variations in drug resistance rates. Drug resistance rate obtained from NAUTH (81.80%) is higher than that reported by [15] that showed 46.1% resistance rate to at least one drug in a study conducted in Nnewi. This indicates an increasing trend in drug resistance which could be as a result of selective compliance to treatment and default among clients [23, 26]. However, since previous rate of drug resistance at St Patrick’s Hospital could not be assessed, a comparison could not be made.

Among the socio-demographic factors assessed in this study, female gender was significantly associated with drug resistance. The resistance rate for the females was higher than that for the males. This agrees with a study conducted in Georgia which revealed that women were more at risk of drug resistance compared to men [27, 24]. The role of women as care givers predispose them to developing drug resistance as they have longer contact at home with sick relatives. Also because of cultural restrictions, women are educationally disadvantaged. Women due to ignorance may not fully understand the importance of adherence to therapy. In a study to access factors contributing to treatment adherence in Zambia, (39.1%) of the females compared with (33.9%) of the males stopped taking their medication after 2 months. Most of the male TB patients were older and more educated than the female TB patients [28]. Gender as a significant demographic factor for drug resistance in this study agrees with the report of [23].

The significant association of smoking with drug resistance in this study has been collaborated in other studies. According to [29] smoking among other life style habits has been associated with development of drug resistance. This agrees with [30] who reported that poor treatment outcomes were higher in smokers in a study in Georgia. [31] Described an association between drug resistance and smoking or tobacco use in some cases of drug resistant TB. It was explained that cigarette smoke contains mutagenic chemicals; and smoking and environmental pollutants could also alter the redox balance, in turn affecting the mutation rate. Significant association of a history of TB treatment in this study with drug resistance agrees with several published articles. [15] Reported that previous TB treatment was a risk factor for MDR-TB and [20] in a study in South-west Nigeria reported that the most significant factor associated with drug resistance was a history of previous anti-TB treatment. Also, [21] reported that previous TB treatment was significant for drug resistance. [22] In a study in Karnataka region, India, showed that past history of pulmonary TB was statistically associated with development of drug resistance. [32] Reported also that previous history of TB treatment among other risk factors was independently associated with high risk of resistance to any first-line anti-TB drug. Delayed recognition of drug resistance, inappropriate chemotherapy regimens, inadequate or irregular drug supply, poor compliance by patients, malabsorbtion of one or more drugs, and sequestered disease (in which differential penetration of anti TB drugs may lead to mono-therapy), have been reported as reasons for development of drug resistance in previously treated patients [26].

Primary drug resistance in this study was high and this is in conformity with [33] who reported resistance rate of (84.6%) in a study in Tanzania. Also [15] reported that a higher proportion of drug resistance was seen among new TB cases than among previously treated cases. [24] Also showed a primary drug resistance rate of (67.12%) in Georgia. However [34, 22, 32], reported lower rates of 10.3%, 9.1% and 15.3% respectively in individual studies done in Uganda, India and Ethiopia respectively. Primary drug resistance occurs when drug resistant bacilli are transmitted to other people [21]. The high rate of primary drug resistance in this study indicates the high proportion of the population harbouring resistant strains. Studies have shown that transmission of drug resistant strains (i.e. primary drug resistance) rather than amplification from susceptible strains (acquisition of resistance conferring mutations i.e. acquired resistance) is the dominant source of MDR-TB [35]. In addition, most of the first line anti-TB drugs are available without prescription (over the counter), and there are no effective control of the availability of these drugs outside the National TB control programme, some of these patients may have been treated with some of these first line anti-TB drugs unknowingly. Hence they are not frank cases of primary drug resistance.

Isolates in this study exhibited the highest resistance rate to Isoniazid, and the least to Ethambutol. [36] In a review of first-line anti-tuberculosis drug resistance reported that Isoniazid had the highest resistance in Iran, in over 16-year while Ethambulol had the least. [15] Showed that Isoniazid mono-resistance was highest among the first line anti-TB drugs and similar pattern of drug resistance was also observed by [32]. Isoniazid is one of the main drugs for TB treatment and over the years increasing levels of resistance to Isoniazid might be due to incomplete treatment [37, 36]. The increased prevalence of strains with primary resistance to Isoniazid is a very important indication to estimate the risk of development of MDR TB [26]. In the same vein, a high proportion of isolates in this study exhibited resistance to Rifampicin. Apprehensively, resistance to Rifampicn is the most pressing concern in the TB management because it necessitates very long, expensive and relatively toxic drug schedules and leads to poorer outcomes [35]. It has been shown that patients infected with strains resistant to Rifampicin will experience a higher failure rate with short course of 6 months chemotherapy [36]. Resistant isolates in this study exhibited distinct patterns of drug resistance. Varying patterns of drug resistance was shown by these isolates with respect to site and treatment history. In treatment of MDR-TB, the number of drugs in the regimen depends on the susceptibility pattern, availability of first line agents and extent of disease [38]. The pattern of drug resistance varies from place to place at different periods of time therefore, knowledge of geographic variations is essential for monitoring of antibiotic resistance within a defined population of patients infected with *M. tuberculosis* [36].

From this study, it can be concluded that there is high prevalence of resistance to first line anti-TB drugs, with female gender, smoking and previous TB treatment being significantly associated with development of drug resistance. It was also observed that there is ongoing community transmission of drug resistance as shown by the high proportion of new cases showing drug resistance and that prevalence of MDR-TB is higher than that documented previously. Worthy of note is that primary transmission of MDR-TB is on the increase, therefore, there is need for more pro-active measures to tackle this public health menace.

## ACKNOWLEDGEMENT

We thank the entire staff of Dr Lawrence Henshaw Memorial Hospital (DLHMH) in Calabar, Cross River State for their assistance and for allowing us to make use of there reference laboratory. This research work received no specific grant from any funding agency in public, commercial, or non-for-profit sector.

